# Chromosome-scale assembly of the *Sparassis latifolia* genome obtained using long-read and Hi-C sequencing

**DOI:** 10.1101/2021.01.08.426014

**Authors:** Chi Yang, Lu Ma, Donglai Xiao, Xiaoyu Liu, Xiaoling Jiang, Zhenghe Ying, Yanquan Lin

## Abstract

*Sparassis latifolia* is a valuable edible mushroom cultivated in China. In 2018, our research group reported an incomplete and low quality genome of *S. latifolia* was obtained by Illumina HiSeq 2500 sequencing. These limitations in the available genome have constrained genetic and genomic studies in this mushroom resource. Herein, an updated draft genome sequence of *S. latifolia* was generated by Oxford Nanopore sequencing and the Hi-C technique. A total of 8.24 Gb of Oxford Nanopore long reads representing ~198.08X coverage of the *S. latifolia* genome were generated. Subsequently, a high-quality genome of 41.41 Mb, with scaffold and contig N50 sizes of 3.31 Mb and 1.51 Mb, respectively, was assembled. Hi-C scaffolding of the genome resulted in 12 pseudochromosomes containing 93.56% of the bases in the assembled genome. Genome annotation further revealed that 17.47% of the genome was composed of repetitive sequences. In addition, 13,103 protein-coding genes were predicted, among which 98.72% were functionally annotated. BUSCO assay results further revealed that there were 92.07% complete BUSCOs. The improved chromosome-scale assembly and genome features described here will aid further molecular elucidation of various traits, breeding of *S. latifolia*, and evolutionary studies with related taxa.

**Importance:** *Sparassis latifolia* Y.C. Dai et Z. Wang (*Sparassidaceae, Polyporales, Agaricomycetes*), collections from Asian *Sparassis* (Dai et al., 2006) exhibits diverse biological and pharmacologic activities. To date, total fresh fruit production in Chinese factories is over 20 tons/d. Despite the significant economic and medical value of *S. latifolia*, the assembly level of genome was under chromosome-scale. This study assembles a high-quality chromosome-scale reference genome of *S. latifolia* using Oxford Nanopore sequencing combined with Hi-C (High-through chromosome conformation capture) scaffolding. The improved reference genome will facilitate molecular breeding of *S. latifolia* and advance our understanding of its genetics and evolution.

## 1 Introduction

*Sparassis latifolia* Y.C. Dai et Z. Wang (*Sparassidaceae, Polyporales, Agaricomycetes*), collections from Asian *Sparassis* (Dai et al., 2006) exhibits diverse biological and pharmacologic activities (Thi Nhu Ngoc et al., 2018; Uchida et al., 2019; Wang et al., 2019). *S. latifolia* is the commonly cultivated *Sparassis* species in China (Yang et al., 2017). To date, total fresh fruit production in Chinese factories is over 20 tons/d. Despite the significant economic and medical value of *S. latifolia*, its genetic information remains limited.

In 2018, our group reported that the *S. latifolia* has a size of 48.13 megabases (Mb) and 12,471 predicted genes (Xiao et al., 2018). Based on this genome sequence, we explored the mechanism of light response and primordia formation of *S. latifolia* (Xiao et al., 2017; Yang et al., 2019). The genome of *S. latifolia* was also deposited at the Joint Genome Institute (JGI, project Id: 1105659) and Genebank (PRJNA562364), consiting of 35.66 Mb and 39.32 Mb genome lengths, respectivily. *S. crispa* is another *Sparassis* species with a reported genome size of 39.0 Mb encoding for 13,157 predicted genes (Kiyama et al., 2018). Another report showed the *S. crispa* genome is 40.406 Mb in length, and contains 18,917 predicted contigs. They also revealed that the complete mitochondrial genome of *S. crispa* is 139,253 bp long, containing 47 genes (Bashir et al., 2020). However, the assembly level of all these studies was under chromosome-scale.

The Oxford Nanopore sequencing (ONT) reads the nucleotide sequence by detecting changes in electrical current signals when a DNA molecule is forced through a biological nanopore (Chen et al., 2020a). Compared to the short reads generated by illimina sequencing, the much longer reads produced by ONT span larger genome regions, resulting in more complete assemblies (Jain et al., 2016). For ONT data, the highest contiguity is obtained with a long-read polished assembly, as compared to a hybrid assembly incorporating both the short and long reads. ONT also significantly improves the assembly completeness as compared to the assembly generated using Illumina reads only (Murigneux et al., 2020). Besides, research on signal simulation of nanopore sequences is highly desirable for method developments of nanopore sequencing applications.

This study aimed to assemble a high-quality chromosome-scale reference genome of *S. latifolia* using ONT combined with Hi-C scaffolding. The improved reference genome will facilitate molecular breeding of *S. latifolia* and advance our understanding of its genetics and evolution.

## 2 Methods

### 2.1 DNA preparation and sequencing

The SP-C strain of *S. latifolia* was grown and maintained on potato dextrose agar (PDA) slants and preserved at the Institute of Edible Fungi, Fujian Academy of Agricultural Sciences (Fuzhou, China). Genomic DNA was isolated from the mycelium of *S. latifolia* by the cetyl-trimethyl ammonium bromide (CTAB) method (Biel and Parrish, 1986). The gDNA was then size-selected and sequenced on an Oxford Nanopore PromethION system by BioMarker Technology Co., Ltd. (Beijing, China). Low-quality reads, reads with adapter sequences, and reads shorter than 2000 nt were filtered out before assembly.

### 2.2 Genome assembly

The *S. latifolia* genome was assembled using NECAT v0.01 (Chen et al., 2020b), then polished by Pilon (Walker et al., 2014) with Illumina short reads, to further eliminate Indel and SNP (single nucleotide polymorphism) errors. BUSCO v3 assessment using single-copy orthologous genes was subsequently performed to confirm the quality of the assembled genome (Simão et al., 2015).

### 2.3 Hi-C library construction and assembly of the chromosome

Genomic DNA extraction, library preparation, and sequencing were carried out by Biomarker Technologies, Beijing, China. Hi-C sequencing libraries were constructed and their concentration and insert size detected using Qubit2.0 and Agilent 2100. Samples with high-quality nuclei were subjected to the Hi-C procedure (Yang et al., 2018). The chromatin was digested using *Hin*d III restriction enzyme and ligated together *in situ* after biotinylation. DNA fragments were subsequently enriched via the interaction of biotin and blunt-end ligation and then subjected to Illumina HiSeq sequencing. Clean reads were mapped to the *S. latifolia* genome using BWA (Li and Durbin, 2009) under its default parameters. Paired-end reads were separately mapped to the genome, followed by filtering out of dangling ends, self-annealing sequences, and dumped pairs. Valid paired-end reads of unique, mapped paired-end reads were collected using HiC-Pro (v2.10) (Servant et al., 2015). The order and direction of scaffolds/contigs were clustered into super scaffolds using LACHESIS (Burton et al., 2013), based on the relationships among valid reads.

### 2.4 Genome annotation

Identification and construction of the *de novo* repeat library were performed by LTR_FINDER v1.05 (Xu and Wang, 2007), MITE-Hunter (Han and Wessler, 2010), RepeatScout v1.0.5 (Price et al., 2005), and PILER-DF v2.4 (Edgar and Myers, 2005). PASTEClassifier (Wicker et al., 2007) was used to classify the database then merged with Repbase's (Jurka et al., 2005) database to generate the final repeat sequence. The RepeatMasker v4.0.6 (Chen, 2004) software was finally used to search the known repeat sequences and map them onto the *de novo* repeat libraries. This step was done to identify novel repeat sequences based on the built repeat sequence database.

The combined use of *ab initio* prediction, homology-based prediction, and transcriptome-assisted prediction was used to identify protein-coding genes. For *ab initio* prediction, Augustus (v3.2.3) (Stanke et al., 2006), Geneid (v1.4)(Alioto et al., 2018), Genescan (v1.0) (Burge and Karlin, 1997), GlimmerHMM (v3.04) (Majoros et al., 2004), and SNAP (v2013.11.29) (Korf, 2004) software were employed under their default parameters. The GeMoMa v1.3.1 software (Jens et al., 2016) was used for homologous protein-based prediction. RNA-seq data were mapped to the sponge gourd genome using Hisat2 (v2.0.4) and Stringtie (v1.2.3) (Pertea et al., 2016) for transcriptome-based prediction. Amino acid sequences were predicted from the assemblies using TransDecoder (v2.0) (The Broad Institute, Cambridge, MA, USA). The results were integrated using the EVM (v1.1.1) (Haas et al., 2008) software to predict all genes.

The protein sequences were subsequently aligned to protein databases for gene annotation. The databases included Gene Ontology (GO) (Ashburner et al., 2000), Kyoto Encyclopedia of Genes and Genomes (KEGG) (http://www.genome.jp/kegg/) (Ogata et al., 1999), InterPro (https://www.ebi.ac.uk/interpro/) (Jones et al., 2014), Swiss-Prot (http://www.uniprot.org) (Boeckmann et al., 2003), and TrEMBL (http://www.uniprot.org/) (Boeckmann et al., 2003). Detection of reliable tRNA positions was accomplished by tRNAscan-SE (v2.0.3) (Lowe and Eddy, 1997). Noncoding RNAs (ncRNAs) were predicted through an RFAM (v12.0) (Griffiths-Jones et al., 2005) database search using the Infernal software (v1.0) (Nawrocki and Eddy, 2013) under its default parameters.

### 2.5 Comparative genomics analysis

Putative orthologous genes were constructed from two *S. latifolia* (SP-C strain in this study and CCMJ1100 strain in JGI) and one *S. crispa* (Kiyama et al., 2018). The OrthoMCL (Li et al., 2003) software was used to classify the protein sequences and analyze the gene families. The classification involved a statistical analysis of the gene families unique to each strain, the gene families shared by the strains, and single-copy gene families for each strain. The gene families were functionally annotated in the Pfam database and their Venn diagram constructed based on their statistical results. The PAML (Yang, 2007) software was then used to calculate Ka/Ks ratios of gene pairs in single-copy gene families. Genes with Ka/Ks ratios greater than 1 were set as fast-evolving genes. Evolutionary trees based on single-copy gene families were constructed using the phyML (Guindon et al., 2010) software to study the evolutionary relationships between species. Comparisons of SP-C protein sequences with each reference genome were made through BLAST analysis (Altschul et al., 1997). Nucleic acid level crosstalk between the genomes pairs was then obtained based on the position of the homologous genes on the genome sequence and plotted using MCScanX (Wang et al., 2012).

## 3 Results and discussion

### 3.1 Sequencing and assembly of the genome

The sequencing depth was 199.08X, which yielded 8.86 Gb of genomic data. Removal of adapters yielded 8.24 Gb of clean data and 14.42 kb of subread N50. The assembled genome had 22 scaffolds, and N50 had significantly increased to 3.31 Mb, compared to 472 scaffolds with N50 of 0.46 Mb (Xiao et al., 2018). Assembly statistics are summarized in Table S1. The primary contigs were further filtered using the Pilon program (Walker et al., 2014) with Illumina short reads to improve accuracy. The post-correction genome size was 41,412,529 bp, with a contig N50 of 1,509,579 bp. The average GC content in the corrected genome was 51.51%.

### 3.2 Assessment of genomic integrity

Approximately 98.74% of the Illumina resequencing reads were mapped to the assembly (Table S2). BUSCO assay results further revealed that there were 92.07% complete BUSCOs (Table S3) which indicated that the assembly integrity was adequate.

### 3.3 Hi-C

The Hi-C approach efficiently uses high-throughput sequencing to determine the state of genome folding by measuring the contact frequency between loci pairs (Lieberman-Aiden et al., 2009). Nearly 39.5 million paired-end reads (11.8 Gb) were collected with a GC content of 53.12% and a Q20 ratio of 97.24% (the percentage of clean reads more than 20 bp) (Table S4). Hi-C library quality was assessed based on the read mapping ratio and the content of invalid interaction pairs. A high-quality Hi-C library plays a vital role in increasing the final effective data volume, which directly reflects the quality of Hi-C library construction. Invalid interaction pairs mainly include self-circle ligation, dangling ends, re-ligation, and dumped pairs (Belton et al., 2012; Imakaev et al., 2012; Servant et al., 2015). The ratio of mapped reads and valid interaction pairs was 90.83% and 86.67%, respectively (Table S5 and S6).

Hi-C assembly located 38,744,916 bp genomic sequences, accounting for 93.56% of the total sequence length on the 12 chromosomes. There were 39 corresponding sequences accounting for 79.59% of the entire sequence. The sequences mapped to the chromosome that determined the chromosomal order and direction were 38,744,916 bp long, accounting for 100% of the total length of the sequence. There were 39 corresponding sequences accounting for 100% of the sequence mapped to the chromosome. Detailed sequence distribution of each chromosome is outlined in Table 2.

**Table 1.** Statistics of *Sparassis* taxa genome.

| Feature | *S. latifolia*<br>(this study) | *S. latifolia*<br>(Xiao et al., 2018) | *S. latifolia*<br>(JGI) | *S. crispa*<br>(Kiyama et al., 2018) | *S. crispa*<br>(Bashir et al., 2020) | *S. latifolia*<br>(Sampled in Larch tree) |
| --- | --- | --- | --- | --- | --- | --- |
| Scaffold Length | 41412529 | 48134914 |  |  |  | 39318455 |
| Scaffold Number | 22 | 472 | 184 |  |  |  |
| Contig Number | 49 | 3848 | 184 | 32 | 18917 | 93 |
| GC content (%) | 51.51 | 51.43 |  | 51.4 |  | 51.4 |
| Sequencing read | 199.08 X | 601X | 74.16x |  |  |  |
| Number of | 13103 | 12471 | 12815 | 13157 |  |  |
| Length of the | 41.41 | 48.13 | 35.66 | 39 | 40.41 | 39.32 |
| mitochondrial |  |  | 133796 |  | 139253 |  |
| Sequencing | ONT | Illumina | PacBio | PacBio RSII |  | Oxford |
| Accession | PRJNA68 | PRJNA31856 | 110565 | PRJDB5582 |  | PRJNA562364 |

**Table 2.** Summary of Hi-C-assisted assembly chromosome lengths.

| Group | Sequence Number | Sequence Length (bp) |
| --- | --- | --- |
| Lachesis Group0 | 3 | 1,652,215 |
| Lachesis Group1 | 2 | 1,841,158 |
| Lachesis Group2 | 2 | 1,873,959 |
| Lachesis Group3 | 2 | 3,372,947 |
| Lachesis Group4 | 1 | 4,516,377 |
| Lachesis Group5 | 3 | 3,305,915 |
| Lachesis Group6 | 3 | 3,178,441 |
| Lachesis Group7 | 5 | 3,208,453 |
| Lachesis Group8 | 4 | 2,878,786 |
| Lachesis Group9 | 7 | 5,687,837 |
| Lachesis Group10 | 5 | 4,576,772 |
| Lachesis Group11 | 2 | 2,652,056 |
| Total Sequences Clustered (Ratio %) | 39(79.59) | 38,744,916 (93.56) |
| Total Sequences Ordered and Oriented (Ratio %) | 39(100) | 38,744,916 (100) |

For the Hi-C assembled into the chromosome, the genome was cut into 20 kb bins with equal length. The number of Hi-C read pairs covered between any two bins was then used as the intensity signal of the interaction between the bins to construct a heat map. The heat map (Fig. 1) revealed that the genome was divided into multiple chromosomes. The interaction intensity at the diagonal position within each group was higher than that of the off-diagonal position, indicating that the interaction between adjacent sequences (diagonal position) in the Hi-C assembly was high. However, there was a weak interaction signal between non-adjacent sequences (off-diagonal positions), which is consistent with the principle of Hi-C assisted genome assembly. This finding suggests that the genome assembly effect was good.

**Figure 1.**
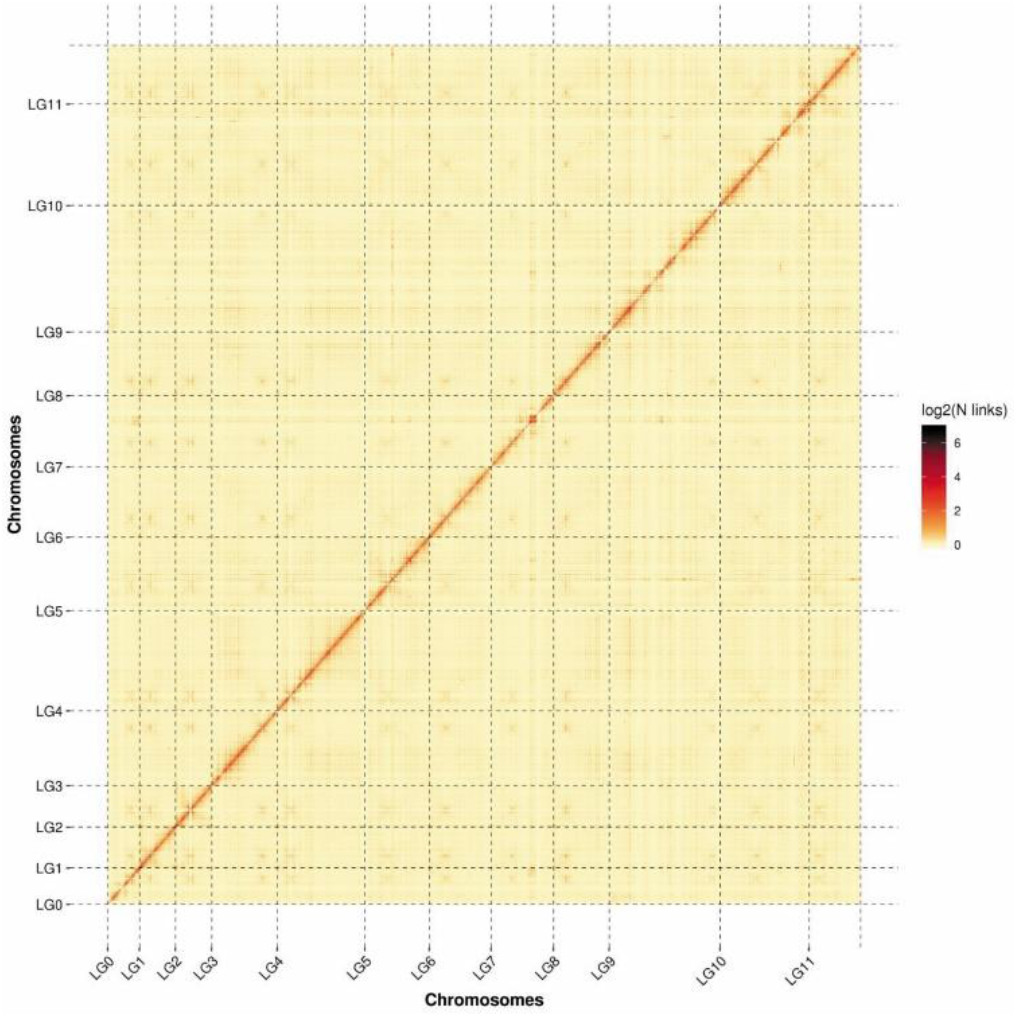
Intensity signal heat map of the Hi-C chromosome. Lchesis Group (LG) means chromosome.

### 3.3 Genome annotation

The *S. latifolia* genome contained 7.23 Mb repetitive sequences that accounted for 17.47% of the genome (Table S7), which was longer than obtained in our previous study (5.19 Mb, 10.79%) (Xiao et al., 2018). Five major types of repeats were detected: class I, class II, potential host gene, SSR, and unknown duplications. Among them, class I comprised the largest proportion (11.11%, total length of 4.60 Mb) followed by the novel repeats (4.54% total length of 1.88 Mb) of the genome.

Annotation was done for 13,103 protein-coding genes in *S. latifolia* (Table S8). Among them, 12,936 (98.72%) were supported by transcriptome data and homolog prediction. The number of protein-coding genes was higher than that of *S. latifolia* previously reported by our group (12,471) (Xiao et al., 2018) and JGI (Project Id: 1105659). However, it was lower than that of *S. crispa* (13,157) (Kiyama et al., 2018). The average length of the predicted genes was 1882.98 bp, while that of their coding sequences was 220.98 bp. In addition, there was an average of 6.2 exons per gene with a length of 238.51 bp per exon. The average intron length was 77.64 bp (Table S9). Functional annotation further revealed that 5,608, 3,341, 7,344, 5,967, 11,187,11,252 and 3,497 genes were annotated to the KOG, GO, Pfam, Swissprot, TrEMBL,nr and KEGG databases, respectively. Cognizant to this, 11,281 (86.09% of the total) genes had at least one hit to the public databases (Table S10). In addition, 126 transfer RNAs, 75 ribosomal RNAs, and 36 other non-coding RNAs were identified in the *S. latifolia* genome (Table S11). There were also 452 identified pseudogenes with premature stop codons or frameshift mutations (Table S12). Nevertheless, this number was significantly higher than previously reported by our group (8 pseudogenes) (Xiao et al., 2018).

### 3.4 Comparison of the genomes of the *Sparassis* taxa

OrthoMCL (Li et al., 2003) was used to classify gene families with single and multiple copies from *Sparassis* taxa, resulting in 909 *S. latifolia* SPC-specific genes (Table 3). *S. latifolia* SPC had more common genes with *S. crispa* SCP (10,083) than with *S. latifolia* CCMJ1100 (8813) (Fig. 2A). However, phylogenetic analyses revealed that *S. latifolia* SPC was more closely related to *S. latifolia* CCMJ1100 (Fig. 2B). Synteny and collinearity analysis between genomes was conducted using MCScanX (Wang et al., 2012) to further characterize the genomic differences between the newly sequenced *S. latifolia* SPC genomes and the other *Sparassis* taxa strains. *S. latifolia* SPC and *S. crispa* SCP genomes were found to be highly collinear (Fig. 2C and 2D). Collinearity analysis further confirmed that the assembled genome was of high quality.

**Table 3.** Statistics of comparison of the genomes of the Sparassis taxa.

| Species Name | Total Gene | Cluster Gene | Total Family | Unique Gene Family | Unique Gene | Ka/Ks>1 |
| --- | --- | --- | --- | --- | --- | --- |
| *S. latifolia* SPC | 13,103 | 12,286 | 10,401 | 42 | 909 |  |
| *S. latifolia* CCMJ1100 | 12,815 | 10,471 | 9,189 | 117 | 2672 | 371 |
| *S. crispa* SCP | 13,157 | 12,103 | 10,309 | 48 | 1164 | 377 |

**Figure 2.**
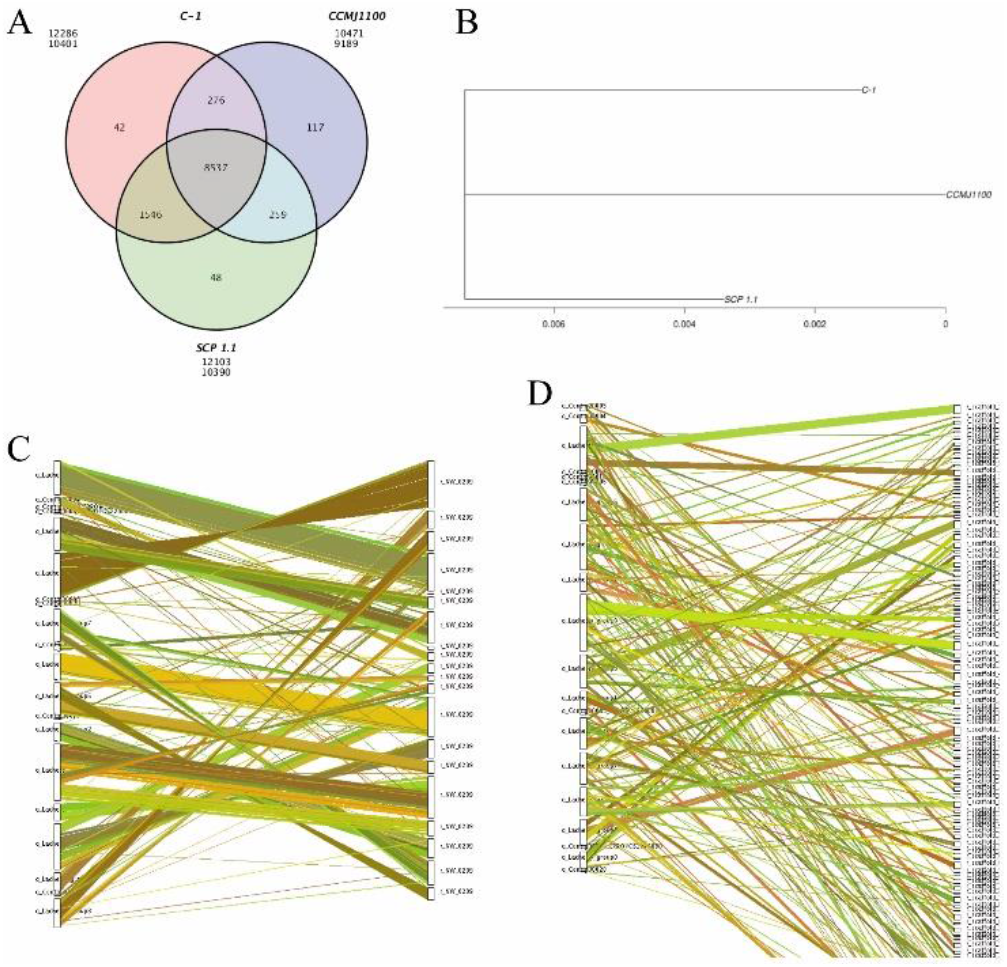
Comparison of the genomes of the *Sparassis* taxa. (A) Gene family annotation Venn diagram. (B) phylogenetic tree. (C-D) Collinearity of *S. latifolia* SPC with *S. crispa* SCP and *S. latifolia* CCMJ1100.

## 4 Conclusions

A 41.41 Mb chromosome-level reference genome of *S. latifolia* was assembled, and its 13,103 protein-coding genes were annotated. The improved assembly and genome features described herein will aid further molecular elucidation of various traits, breeding of *S. latifolia*, and evolutionary studies with related taxa.

## Data availability

Genome assembly was submitted to NCBI under the BioProject accession number PRJNA686158.

## Acknowledgments

This work was supported by the Natural Science Foundation of Fujian province of China (2020J011378), the Special Fund for Scientific Research in the Public Interest of Fujian Province (2020R1035003, 2020R1035005).

## Reference

Alioto, T., Blanco, E., Parra, G., and Guigó, R. (2018). Using geneid to Identify Genes. Curr Protoc Bioinformatics 64(1), e56. doi: 10.1002/cpbi.56.

Altschul, S.F., Madden, T.L., Schaffer, A.A., Zhang, J., Zhang, Z., Miller, W., et al. (1997). Gapped BLAST and PSI-BLAST: a new generation of protein database search programs. Nucleic Acids Res 25(17), 3389–3402. doi: 10.1093/nar/25.17.3389.

Ashburner, M., Ball, C.A., Blake, J.A., Botstein, D., Butler, H., Cherry, J.M., et al. (2000). Gene ontology: tool for the unification of biology. The Gene Ontology Consortium. Nat Genet 25(1), 25–29. doi: 10.1038/75556.

Bashir, K.M.I., Rheu, K.M., Kim, M.-S., and Cho, M.-G. (2020). The complete mitochondrial genome of an edible mushroom, Sparassis crispa. Mitochondrial DNA Part B 5(1).

Belton, J.M., McCord, R.P., Gibcus, J.H., Naumova, N., Zhan, Y., and Dekker, J. (2012). Hi-C: a comprehensive technique to capture the conformation of genomes. Methods 58(3), 268–276. doi: 10.1016/j.ymeth.2012.05.001.

Biel, S.W., and Parrish, F.W. (1986). Isolation of DNA from fungal mycelia and sclerotia without use of density gradient ultracentrifugation. Anal Biochem 154(1), 21–55.

Boeckmann, B., Bairoch, A., Apweiler, R., Blatter, M.C., Estreicher, A., Gasteiger, E., et al. (2003). The SWISS-PROT protein knowledgebase and its supplement TrEMBL in 2003. Nucleic Acids Res 31(1), 365–370. doi: 10.1093/nar/gkg095.

Burge, C., and Karlin, S. (1997). Prediction of complete gene structures in human genomic DNA. J Mol Biol 268(1), 78–94. doi: 10.1006/jmbi.1997.0951.

Burton, J.N., Adey, A., Patwardhan, R.P., Qiu, R., Kitzman, J.O., and Shendure, J. (2013). Chromosome-scale scaffolding of de novo genome assemblies based on chromatin interactions. Nat Biotechnol 31(12), 1119–1125. doi: 10.1038/nbt.2727.

Chen, N. (2004). Using RepeatMasker to identify repetitive elements in genomic sequences. Curr Protoc Bioinformatics Chapter 4, Unit 4.10. doi: 10.1002/0471250953.bi0410s05.

Chen, W., Zhang, P., Song, L., Yang, J., and Han, C. (2020a). Simulation of Nanopore Sequencing Signals Based on BiGRU. Sensors (Basel) 20(24). doi: 10.3390/s20247244.

Chen, Y., Nie, F., Xie, S.-Q., Zheng, Y.-F., Bray, T., Dai, Q., et al. (2020b). Fast and accurate assembly of Nanopore reads via progressive error correction and adaptive read selection. bioRxiv, 2020.2002.2001.930107. doi: 10.1101/2020.02.01.930107.

Dai, Y.C., Wang, Z., Binder, M., and Hibbett, D.S. (2006). Phylogeny and a new species of Sparassis (Polyporales, Basidiomycota): evidence from mitochondrial atp6, nuclear rDNA and rpb2 genes. Mycologia 98(4), 584–592.

Edgar, R.C., and Myers, E.W. (2005). PILER: identification and classification of genomic repeats. Bioinformatics 21 Suppl 1, i152–158. doi: 10.1093/bioinformatics/bti1003.

Griffiths-Jones, S., Moxon, S., Marshall, M., Khanna, A., Eddy, S.R., and Bateman, A. (2005). Rfam: annotating non-coding RNAs in complete genomes. Nucleic Acids Res 33(Database issue), D121–124. doi: 10.1093/nar/gki081.

Guindon, S., Dufayard, J.F., Lefort, V., Anisimova, M., Hordijk, W., and Gascuel, O. (2010). New algorithms and methods to estimate maximum-likelihood phylogenies: assessing the performance of PhyML 3.0. Syst Biol 59(3), 307–321. doi: 10.1093/sysbio/syq010.

Haas, B.J., Salzberg, S.L., Zhu, W., Pertea, M., Allen, J.E., Orvis, J., et al. (2008). Automated eukaryotic gene structure annotation using EVidenceModeler and the Program to Assemble Spliced Alignments. Genome Biol 9(1), R7. doi: 10.1186/gb-2008-9-1-r7.

Han, Y., and Wessler, S.R. (2010). MITE-Hunter: a program for discovering miniature inverted-repeat transposable elements from genomic sequences. Nucleic Acids Res 38(22), e199. doi: 10.1093/nar/gkq862.

Imakaev, M., Fudenberg, G., McCord, R.P., Naumova, N., Goloborodko, A., Lajoie, B.R., et al. (2012). Iterative correction of Hi-C data reveals hallmarks of chromosome organization. Nat Methods 9(10), 999–1003. doi: 10.1038/nmeth.2148.

Jain, M., Olsen, H.E., Paten, B., and Akeson, M. (2016). The Oxford Nanopore MinION: delivery of nanopore sequencing to the genomics community. Genome Biol 17(1), 239. doi: 10.1186/s13059-016-1103-0.

Jens, K., Michael, W., Erickson, J.L., Schattat, M.H., Jan, G., and Frank, H. (2016). Using intron position conservation for homology-based gene prediction. Nuclc Acids Research (9), e89–e89.

Jones, P., Binns, D., Chang, H.Y., Fraser, M., Li, W., McAnulla, C., et al. (2014). InterProScan 5: genome-scale protein function classification. Bioinformatics 30(9), 1236–1240. doi: 10.1093/bioinformatics/btu031.

Jurka, J., Kapitonov, V.V., Pavlicek, A., Klonowski, P., Kohany, O., and Walichiewicz, J. (2005). Repbase Update, a database of eukaryotic repetitive elements. Cytogenet Genome Res 110(1-4), 462–467. doi: 10.1159/000084979.

Kiyama, R., Furutani, Y., Kawaguchi, K., and Nakanishi, T. (2018). Genome sequence of the cauliflower mushroom Sparassis crispa (Hanabiratake) and its association with beneficial usage. Sci Rep 8(1), 16053. doi: 10.1038/s41598-018-34415-6.

Korf, I. (2004). Gene finding in novel genomes. BMC Bioinformatics 5, 59. doi: 10.1186/1471-2105-5-59.

Li, H., and Durbin, R. (2009). Fast and accurate short read alignment with Burrows-Wheeler transform. Bioinformatics 25(14), 1754–1760. doi: 10.1093/bioinformatics/btp324.

Li, L., Stoeckert, C.J.Jr., and Roos, D.S. (2003). OrthoMCL: identification of ortholog groups for eukaryotic genomes. Genome Res 13(9), 2178–2189. doi: 10.1101/gr.1224503.

Lieberman-Aiden, E., van Berkum, N.L., Williams, L., Imakaev, M., Ragoczy, T., Telling, A., et al. (2009). Comprehensive mapping of long-range interactions reveals folding principles of the human genome. Science 326(5950), 289–293. doi: 10.1126/science.1181369.

Lowe, T.M., and Eddy, S.R. (1997). tRNAscan-SE: a program for improved detection of transfer RNA genes in genomic sequence. Nucleic Acids Res 25(5), 955–964. doi: 10.1093/nar/25.5.955.

Majoros, W.H., Pertea, M., and Salzberg, S.L. (2004). TigrScan and GlimmerHMM: two open source ab initio eukaryotic gene-finders. Bioinformatics 20(16), 2878–2879. doi: 10.1093/bioinformatics/bth315.

Murigneux, V., Rai, S.K., Furtado, A., Bruxner, T.J.C., Tian, W., Harliwong, I., et al. (2020). Comparison of long-read methods for sequencing and assembly of a plant genome. Gigascience 9(12). doi: 10.1093/gigascience/giaa146.

Nawrocki, E.P., and Eddy, S.R. (2013). Infernal 1.1: 100-fold faster RNA homology searches. Bioinformatics 29(22), 2933–2935. doi: 10.1093/bioinformatics/btt509.

Ogata, H., Goto, S., Sato, K., Fujibuchi, W., Bono, H., and Kanehisa, M. (1999). KEGG: Kyoto Encyclopedia of Genes and Genomes. Nucleic Acids Res 27(1), 29–34. doi: 10.1093/nar/27.1.29.

Pertea, M., Kim, D., Pertea, G.M., Leek, J.T., and Salzberg, S.L. (2016). Transcript-level expression analysis of RNA-seq experiments with HISAT, StringTie and Ballgown. Nat Protoc 11(9), 1650–1667. doi: 10.1038/nprot.2016.095.

Price, A.L., Jones, N.C., and Pevzner, P.A. (2005). De novo identification of repeat families in large genomes. Bioinformatics 21 Suppl 1, i351–358. doi: 10.1093/bioinformatics/bti1018.

Servant, N., Varoquaux, N., Lajoie, B.R., Viara, E., Chen, C.J., Vert, J.P., et al. (2015). HiC-Pro: an optimized and flexible pipeline for Hi-C data processing. Genome Biol 16, 259. doi: 10.1186/s13059-015-0831-x.

Simão, F.A., Waterhouse, R.M., Panagiotis, I., Kriventseva, E.V., and Zdobnov, E.M. (2015). BUSCO: assessing genome assembly and annotation completeness with single-copy orthologs. Bioinformatics (19), 3210–3212.

Stanke, M., Keller, O., Gunduz, I., Hayes, A., Waack, S., and Morgenstern, B. (2006). AUGUSTUS: ab initio prediction of alternative transcripts. Nucleic Acids Res 34(Web Server issue), W435–439. doi: 10.1093/nar/gkl200.

Thi Nhu Ngoc, L., Oh, Y.K., Lee, Y.J., and Lee, Y.C. (2018). Effects of Sparassis crispa in Medical Therapeutics: A Systematic Review and Meta-Analysis of Randomized Controlled Trials. Int J Mol Sci 19(5), 1487. doi: 10.3390/ijms19051487.

Uchida, M., Horii, N., Hasegawa, N., Oyanagi, E., Yano, H., and Iemitsu, M. (2019). Sparassis crispa Intake Improves the Reduced Lipopolysaccharide-Induced TNF-alpha Production That Occurs upon Exhaustive Exercise in Mice. Nutrients 11(9). doi: 10.3390/nu11092049.

Walker, B.J., Abeel, T., Shea, T., Priest, M., Abouelliel, A., Sakthikumar, S., et al. (2014). Pilon: an integrated tool for comprehensive microbial variant detection and genome assembly improvement. PLoS One 9(11), e112963. doi: 10.1371/journal.pone.0112963.

Wang, Y., Tang, H., Debarry, J.D., Tan, X., Li, J., Wang, X., et al. (2012). MCScanX: a toolkit for detection and evolutionary analysis of gene synteny and collinearity. Nucleic Acids Res 40(7), e49. doi: 10.1093/nar/gkr1293.

Wang, Z., Liu, J., Zhong, X., Li, J., Wang, X., Ji, L., et al. (2019). Rapid Characterization of Chemical Components in Edible Mushroom Sparassis crispa by UPLC-Orbitrap MS Analysis and Potential Inhibitory Effects on Allergic Rhinitis. Molecules 24(16). doi: 10.3390/molecules24163014.

Wicker, T., Sabot, F., Hua-Van, A., Bennetzen, J.L., Capy, P., Chalhoub, B., et al. (2007). A unified classification system for eukaryotic transposable elements. Nat Rev Genet 8(12), 973–982. doi: 10.1038/nrg2165.

Xiao, D., Zhang, D., Ma, L., Wang, H., and Lin, Y. (2017). Preliminary study on differentially expressed genes of Sparassis latifolia under light inducing. Edible Fungi of China 36(5), 60–63. (in Chinese). doi: 10.13629/j.cnki.53-1054.2017.05.014.

Xiao, D.L., Ma, L., Yang, C., Ying, Z.H., Jiang, X.L., and Lin, Y.Q. (2018). De Novo Sequencing of a Sparassis latifolia Genome and Its Associated Comparative Analyses. Can J Infect Dis and Med 2018, 12. doi: 10.1155/2018/1857170. eCollection 2018.

Xu, Z., and Wang, H. (2007). LTR_FINDER: an efficient tool for the prediction of full-length LTR retrotransposons. Nucleic Acids Res 35(Web Server issue), W265–268. doi: 10.1093/nar/gkm286.

Yang, C., Ma, L., Xiao, D., Ying, Z., Jiang, X., and Lin, Y. (2019). Integration of ATAC-Seq and RNA-Seq Identifies Key Genes in Light-Induced Primordia Formation of Sparassis latifolia. Int J Mol Sci 21(1), 185. doi: 10.3390/ijms21010185.

Yang, C., Ma, L., Ying, Z.H., Jiang, X.L., and Lin, Y.Q. (2017). Sequence Analysis and Expression of a Blue-light Photoreceptor Gene, Slwc-1 from the Cauliflower Mushroom Sparassis latifolia. Current Microbiology 74(4), 469–475. doi: 10.1007/s00284-017-1218-x.

Yang, X., Yue, Y., Li, H., Ding, W., Chen, G., Shi, T., et al. (2018). The chromosome-level quality genome provides insights into the evolution of the biosynthesis genes for aroma compounds of Osmanthus fragrans. Horticulture research 5, 72–72. doi: 10.1038/s41438-018-0108-0.

Yang, Z. (2007). PAML 4: phylogenetic analysis by maximum likelihood. Mol Biol Evol 24(8), 1586–1591. doi: 10.1093/molbev/msm088.

